# Dynamic reconfiguration of functional subgraphs after musical training in young adults

**DOI:** 10.1101/639856

**Authors:** Qiongling Li, Xuetong Wang, Shaoyi Wang, Yongqi Xie, Yachao Xie, Shuyu Li

## Abstract

The human brain works in a form of network architecture in which dynamic modules and subgraphs were considered to enable efficient information communication supporting diverse brain functions from fixed anatomy. Previous study demonstrated musical training induced flexible node assignment changes of visual and auditory systems. However, how the dynamic subgraphs change with musical training still remains largely unknown. Here, 29 novices healthy young adults who received 24-week piano training, and another 27 novices without any intervention were scanned at three time points—before and after musical training, and 12 weeks after training. We used nonnegative matrix factorization to identify a set of subgraphs and their corresponding time-dependent coefficients from a concatenated functional network of all subjects in sliding time windows. The energy and entropy of the time-dependent coefficients were computed to quantify the subgraph’s dynamic changes in expression. The musical training group showed significantly increased energy of time-dependent coefficients of 3 subgraphs after training. Furthermore, one of the subgraphs, comprised of primary functional systems and cingulo-opercular task control and salience systems, showed significantly changed entropy in the training group after training. Our results suggest that interaction of functional systems undergoes significant changes in their fine-scale dynamic after a period of musical training.

**Author Summary:** We designed a longitudinal experiment to investigate the musical training induced dynamic subgraph changes in 29 novice healthy young adults before and after musical training compared with another 27 novice participants who were evaluated longitudinal but without any intervention. The nonnegative matrix factorization was employed to decompose the constructed dynamic functional connectivity matrix to a set of subgraphs and their corresponding time-dependent coefficients. We found that functional systems interacted closely with each other during transient process, and the musical training group showed significantly increased energy and entropy of time-dependent coefficients after training when compared with the control group. The present study suggests that musical training could induce the reconfiguration of functional subgraphs in young adults.

## Introduction

The human brain works in a form of network architecture in which dynamic modules and subgraphs were considered to enable efficient information communication supporting diverse brain functions from fixed anatomy [1]. Network neuroscience provides tools to help reveal network structure and processes that support integrated brain functions across multiple spatial and temporal scales [2]. In network neuroscience, greatly interconnected nodes known as modules can reconfigure over time when healthy human participants engage in motor skill learning [3], executive cognition tasks [4] and musical training [5]. Musical performance is one of the most complex skills, like instrument performance involving musical notations sight-reading, hand movements, auditory feedback and higher-order cognition mediation, which needs continuous and dynamic information integration of multiple functional systems [6–8]. Previous studies have reported that musical training is related with behavioral, structural, and functional changes on time scales ranging from days to years [9–12].In our previous work, we used sliding window to construct the dynamic functional brain network and quantitatively evaluated the dynamic statistics of the 13 well-known functional systems by identifying putative functional modules in the brain network [5]. We found more flexible node (brain region) assignment changes of visual and auditory systems in the musical training group after musical training. It has been proven that not only the nodes assignment changes during tasks or at resting state, but also the clusters, also known as subgraphs, whose strengths or weights vary together, were dynamic in brain functional network. More importantly, the switching behaviour of the subgraph was considered to be associated with the transience of brain states during development [13] and a key substrate of cognitive control process and behaviour [14]. Di et al [15] demonstrated that the resting-state connectivity variations were associated with the low or high intrinsic activities of specific networks which may reflect the changes in mental state. Musical performance strongly depends on the integration of brain functional systems, while how the dynamic subgraphs change with the musical training still remains largely unknown. The answer is important to reveal the nature of training-induced plasticity and to provide new evidence regarding the functional architecture of the brain network by exploring the dynamic changes of the functional subgraphs before and after musical training in young adults.

Nonnegative matrix factorization (NMF) is an unsupervised machine-learning method [16] to factorizes a matrix into two matrices: features matrix and coefficients matrix, with the property that all three matrices have no negative elements. NMF has been widely used in a variety of domains, such as image processing [17], natural language processing [18], facial and speech recognition [16, 19], and computational biology [20]. When NMF is used for spatially and temporally overlapping subgraph detection in neuroimage, the functional connectivity matrix, concatenated from all subjects’ functional networks in sliding time windows, can be decomposed into a matrix of subgraphs and a matrix of time-dependent coefficients that quantify the level of expression in each time window for each subgraph [13]. Other coactiviation pattern driven approaches, such as principal component analysis (PCA) [21] or independent component analysis (ICA) [22], may yield positive or negative subgraph interactions and time-dependent expression coefficients. NMF enforces nonnegativity giving rise to the nonnegative combination of basis subgraphs and time-dependent expression coefficients, which eases the neurophysiological interpretability of the expressed functional subgraphs over time. Previous studies have shown changes in dynamics of subgraphs over development [13] and cognitive control behaviour [14] by using NMF. This computational tool allows us to track how set of subgraphs are dynamically expressed during experimentally modulated changes in musical training model.

In this study, we applied the NMF to decompose the constructed dynamic functional connectivity matrix to a set of subgraphs and their corresponding time-dependent expression coefficients as shown in Fig 1. The dynamic functional connectivity was constructed from the resting state fMRI of 29 novice healthy young adults in the training group and 27 matched young adults in the control group scanning from 3 time points. Participants in the training group received 24 weeks piano training while participants in the control group without any training. We compared the differences of the subgraph expression between the two groups after and before musical training. We hypothesized that participants in the training group would exhibit more flexible dynamic subgraph expression after musical training compared with the controls without any intervention.

**Fig 1.**
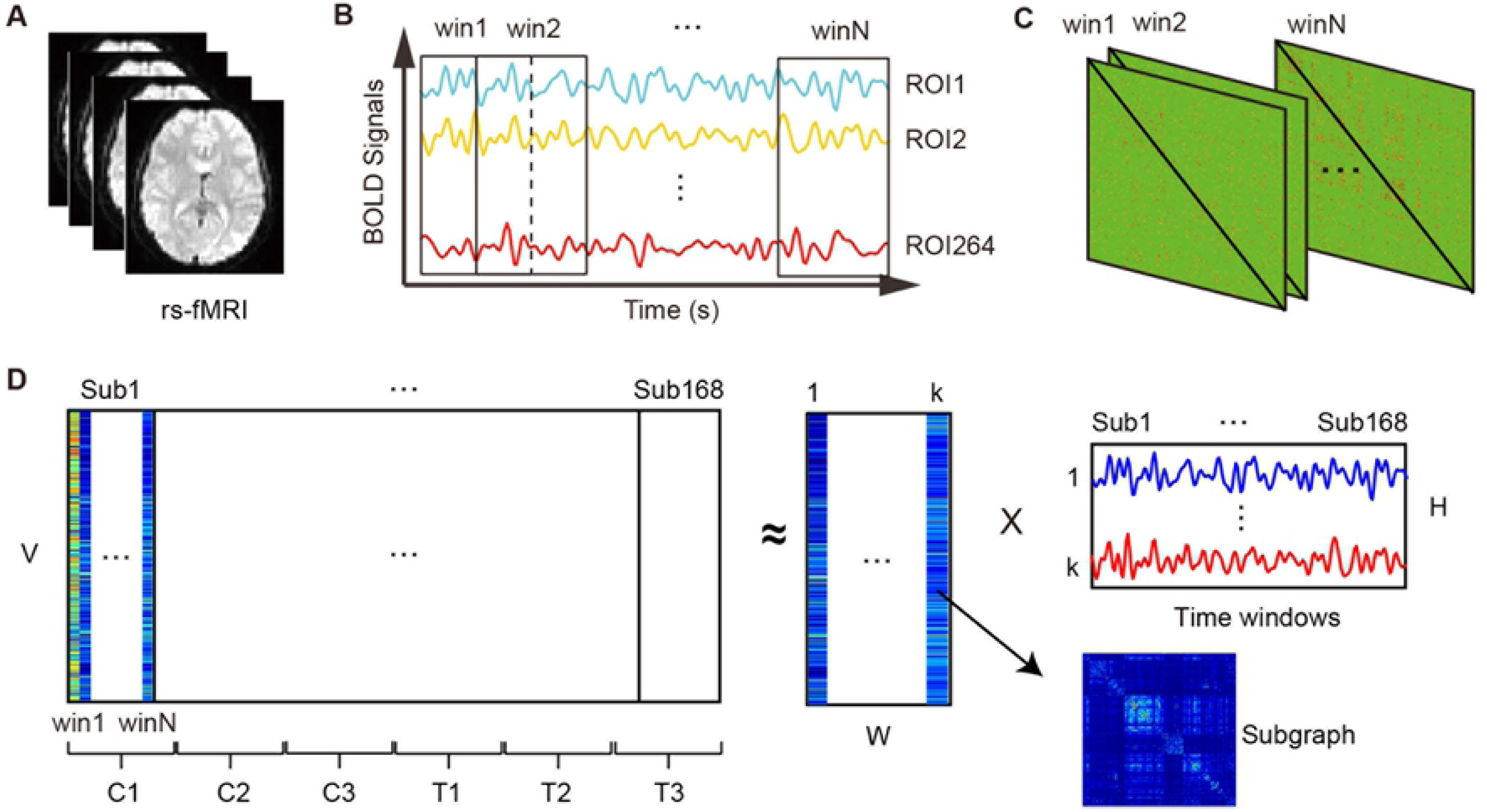
Flowchart of the dynamic functional connectivity expression. (**A**) The resting-state fMRI data were all preprocessed for each subject in each scanning session. **(B)** The average time series for each of the 264 ROIs were divided into 141 time windows Sliding time windows were used to divide the average time series for each of the 264 ROIs into 141 time windows. **(C)** Wavelet coherence was computed between each pair of regional time series for each time window to obtain multiple functional networks for each subject in each scanning session. **(D)** The upper triangle matrix of functional network in each time window for each subject in each scanning session were extracted and unfolded into a vector to obtain a concatenated matrix through time windows for one subject and then through all subjects and scanning sessions (left). Then, a nonnegative matrix factorization method was used to decompose the concatenated matrix into nonnegative matrix W of a set of subgraphs and H of time-dependent coefficients which quantify the level of expression in each time window for each subgraph (right). Notes: C1, control group at time point 1; C2, control group at time point 2; C3, control group at time point 3; T1, training group at time point 1; T2, training group at time point 2; T3, training group at time point 3.

## Results

### Participants and demographics

There were no significant differences in gender, age, education, Beck Depression Inventory (BDI), IQ and Advanced Measures of Music Audiation (AMMA) scores (𝑝 > 0.05) between the control group (age: 23.33 ± 1.39 ys, education: 16.70 ± 1.26 ys, BDI: 5.15 ± 3.87, AMMA: 51.93 ± 15.31) and the training group (age: 23.10 ± 1.37 ys, education: 16.59 ± 1.09 ys, BDI: 4.71 ± 3.62, AMMA: 57.28 ± 11.08) at baseline. And no significant differences were found in the behavioural tests, including digit span, digit symbol, block design, and the trail-making tests (parts A and B) between both groups at baseline. A mixed ANOVA was performed to test the interaction of the group over time, while no significant interaction effects of group over time were found in any of these behavioural tests.

### Subgraphs and their expression coefficients

As shown in Fig 2A, the reconstruction error (residual sum of squares, RSS) is more dependent on the number of subgraphs 𝑘 and less sensitive to changes in the parameter 𝛽, which means robust results of sparse time-dependent coefficients. Parameter 𝛼, which depends on the range and level of sparsity of the expression coefficients 𝐻, was set to be the square of the maximal element in 𝑉 to regulate the magnitude of connection strengths in the subgraphs 𝑊. We observe that the RSS decreases with increasing 𝑘 showing in Figure 2B. Based on the firs-order difference of RSS versus 𝑘, we selected 𝑘 = 10, as steadily increasing derivative RSS when *k* lager than 10. Then, 𝛽 = 10^−1^ was selected for considering the minimum of the RSS against 𝛽 when 𝑘 = 10. After selecting the two parameters, the NMF decomposition was performed to obtain a set of 10 subgraphs and their corresponding time-dependent coefficients for each individual. The obtained 10 subgraphs and the average time-dependent coefficients for both groups at each time point were presented in Fig 3. We can observe that the subgraphs captured varying interactions among brain regions. Some subgraphs showed distributed interactions across the entire network, while other subgraphs showed relatively local interactions, suggesting a complex landscape of the brain.

**Fig 2.**
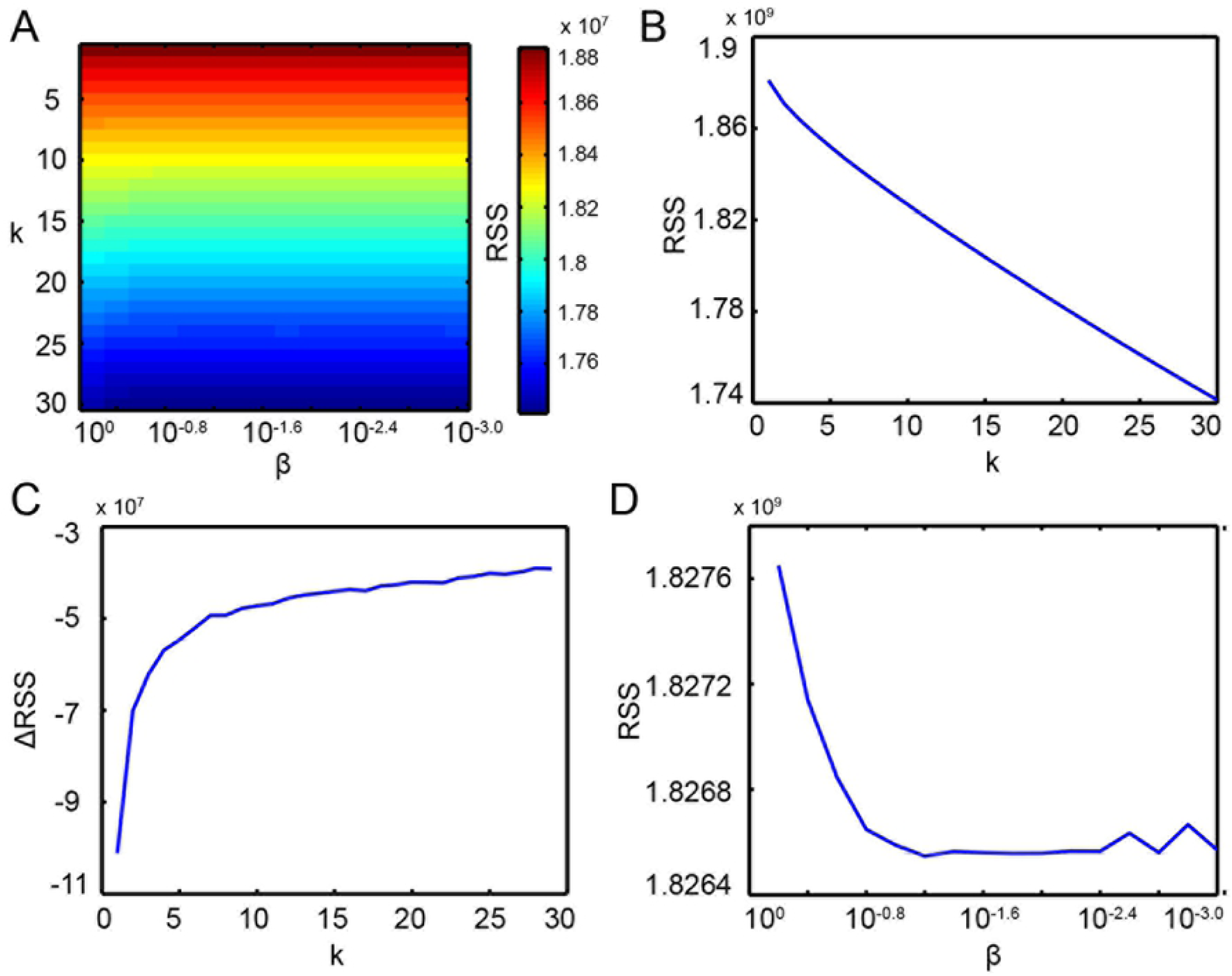
Optimal parameters selection. **(A)** The grid residual sum of squares (RSS) was computed in the range of 𝑘 = [1,⋯,30] and 𝛽 = [10^0^,⋯,10 ^−3^]. **(B)** The averaged RSS across 𝛽 𝛽 versus the number of subgraphs 𝑘 was plotted showing an decrease RSS with increasing 𝑘. **(C)** The first-order difference of RSS versus 𝑘 was plotted and 𝑘 = 10 was selected as steadily increasing derivative RSS when 𝑘 lager than 10. **(D)** The RSS was plotted against 𝛽 𝛽 for 𝑘 = 10 and 𝛽 = 10 ^−1^ was selected to be the first minimum of the steadily decreasing RSS.

**Fig 3.**
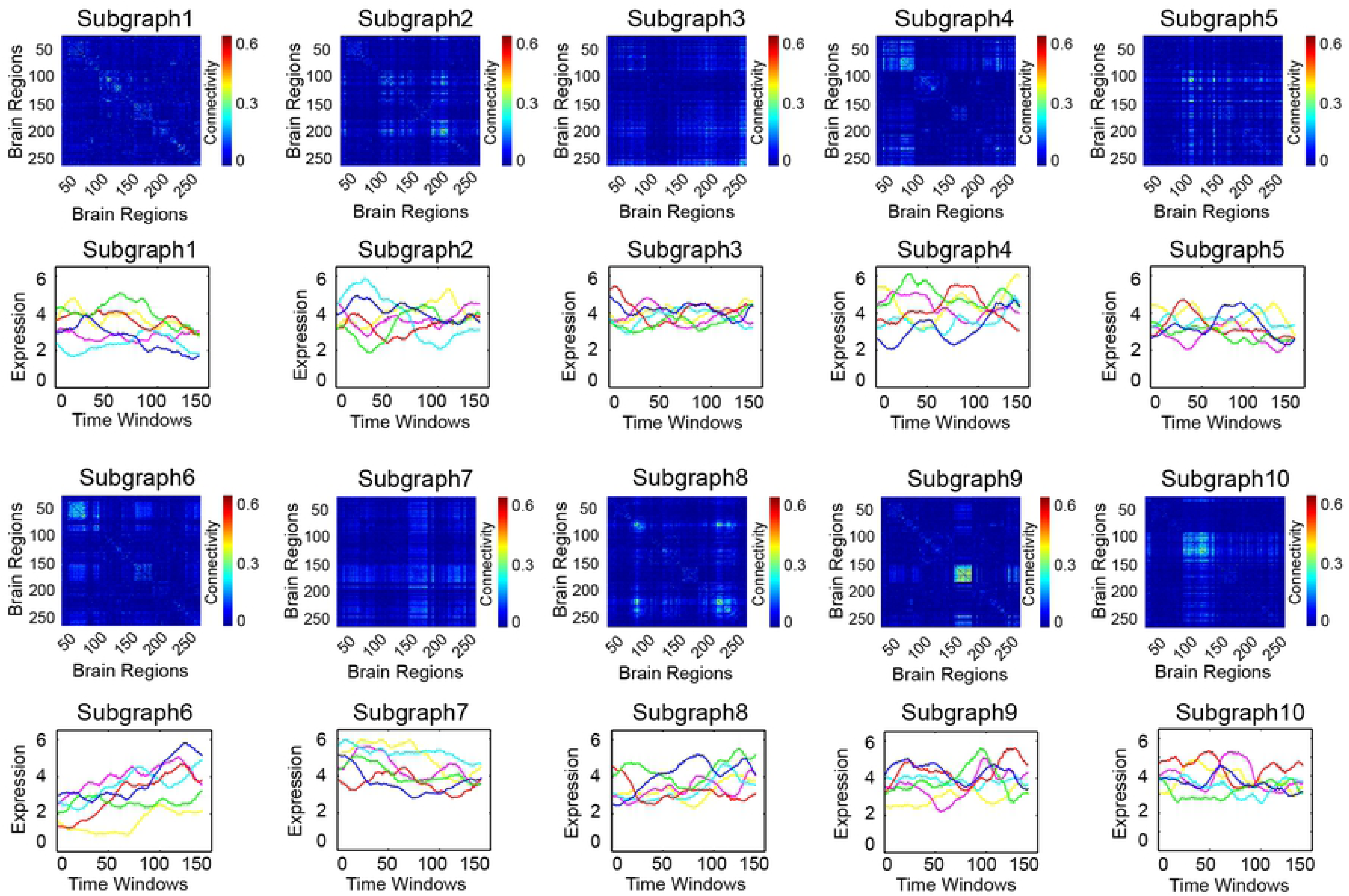
Nonnegative matrix factorization of functional connectivity matrix. A set of 10 subgraphs and their corresponding average time-dependent coefficients for both groups at each time point were obtained from applying nonnegative matrix factorization to the concatenated functional connectivity matrix. Notes: C1, control group at time point 1; C2, control group at time point 2; C3, control group at time point 3; T1, training group at time point 1; T2, training group at time point 2; T3, training group at time point 3.

### Significantly expressed functional systems

Permutation test with 95% confidence interval threshold was performed to detect significantly expressed functional systems for the obtained 10 subgraphs. As presented in Fig 4, black circles indicate significantly expressed functional systems and these systems were mapped into a brain surface which were differentially distributed in each subgraph. All subgraphs were not mapped into individual functional systems in one-to-one manner, which suggested that functional systems do not function as distinct entities over short timescales. Instead, several functional systems were presented during transient process, which together produce a complex brain dynamics landscape that support cognition. We observed that almost all functional systems were expressed in either of the 10 subgraphs except the ventral attention and dorsal attention. The default mode presented a broad expression in all subgraphs, as well as the primary functional systems, such as the sensory-motor, auditory and visual systems. However, functional systems related to higher-order cognition tended to less expressed and usually co-activated with primary functional systems.

**Fig 4.**
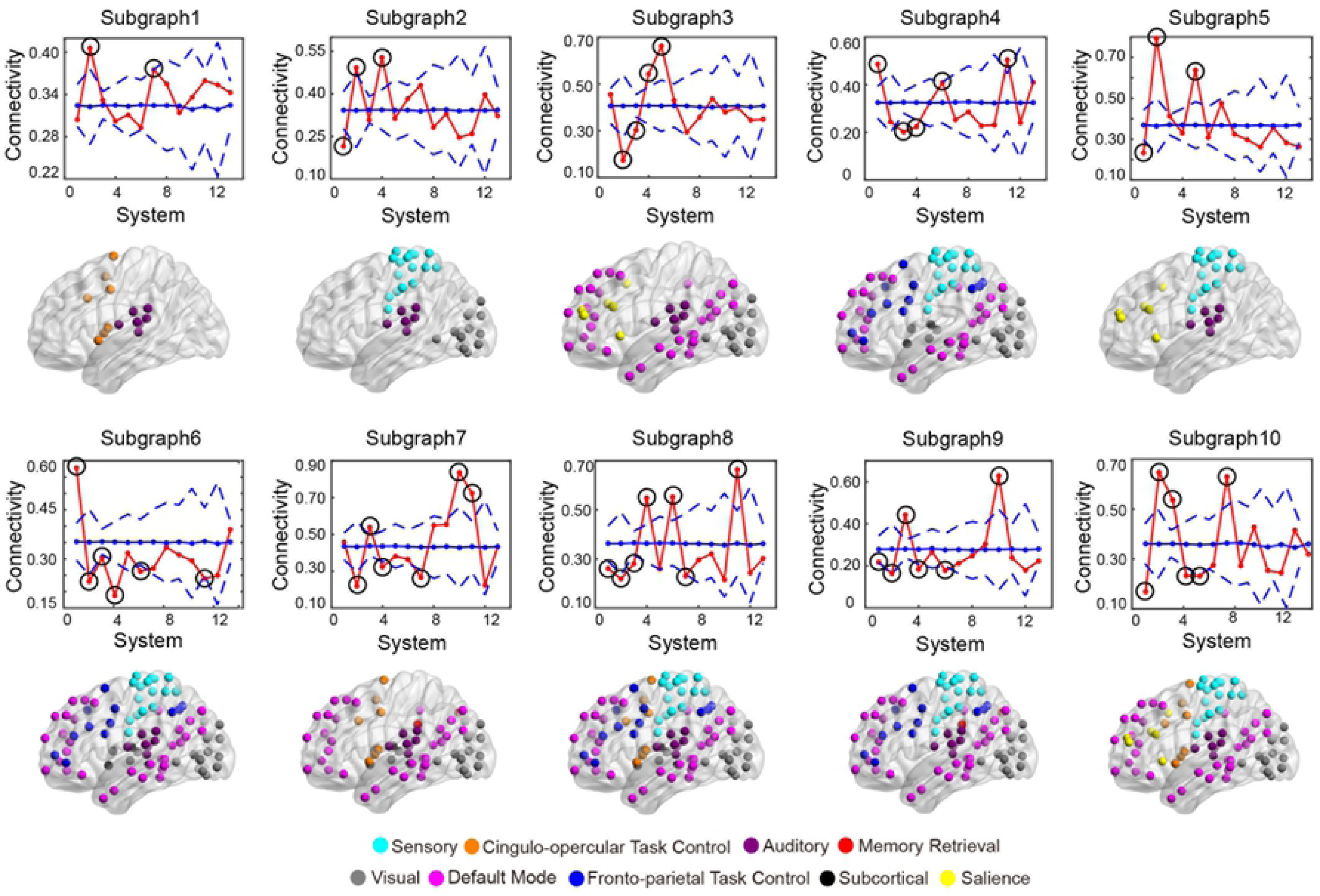
Significantly expressed functional systems. Permutation tests were performed to determine the significantly expressed functional systems. Functional system labels were randomly permuted 1000 times and the mean connectivity in each system was computed to contrast the null distribution. The blue lines represent the mean values (solid lines) and 95% confidence intervals (dash lines) of the mean connectivity for each functional system in the null distribution. The red lines represent the real mean connectivity of each functional system. Those with black circles indicate significantly expressed functional systems whose mean connectivity strength was above the 95% confidence interval threshold. Significantly expressed functional systems in each subgraph were mapped into a brain surface showing a differentially distribution across subgraphs.

### Musical training induced subgraphs differences in expression and dynamics

The energy and entropy of each subgraph’s corresponding time-dependent coefficients were computed and the ANOVA was used to detect the interaction effect of group over time for both quantitative measurements. For energy analyses, significant interaction effects were found in time-dependent coefficients of subgraph 4 (𝐹_2,102_ = 3.257, 𝑝 = 0.043), subgraph 6 (𝐹_2,102_ = 3.639, 𝑝 = 0.030) and subgraph 10 (𝐹_2,102_ = 3.567, 𝑝 = 0.032). Post-hoc comparisons found that participants showed increased energy for all the three subgraphs when comparing Tp2 with Tp1 (subgraph 4, 𝑝 = 0.035; subgraph 6, 𝑝 = 0.005; subgraph 10, 𝑝 = 0.006) in the training group as shown in Fig 5. When comparing Tp3 with Tp2, only subgraph 4 (𝑝 = 0.020) and subgraph 10 (𝑝 = 0.003) showed significant decrease in the training group. Significantly expressed functional systems in subgraph 4 included sensory, default mode, visual, and fronto-parietal task control systems. Subgraph 6 included sensory, default mode, visual, fronto-parietal task control, and auditory systems. And the subgraph 10 included sensory, default mode, visual, auditory, cingulo-opercular task control, and salience systems. For entropy analyses, significant interaction effects were only found in subgraph 10 (𝐹_2,102_ = 7.943, 𝑝 = 0.001) with post hoc pairwise comparisons showing increased entropy when comparing Tp2 with Tp1 (𝑝 < 0.001) and decreased entropy when comparing Tp3 with Tp2 (𝑝 = 0.001) in the training group as shown in Fig 6. All these significant changes were Bonferroni corrected for multiple comparison. Besides, these significant changes showed no correlation with the practice time in the training group.

**Fig 5.**
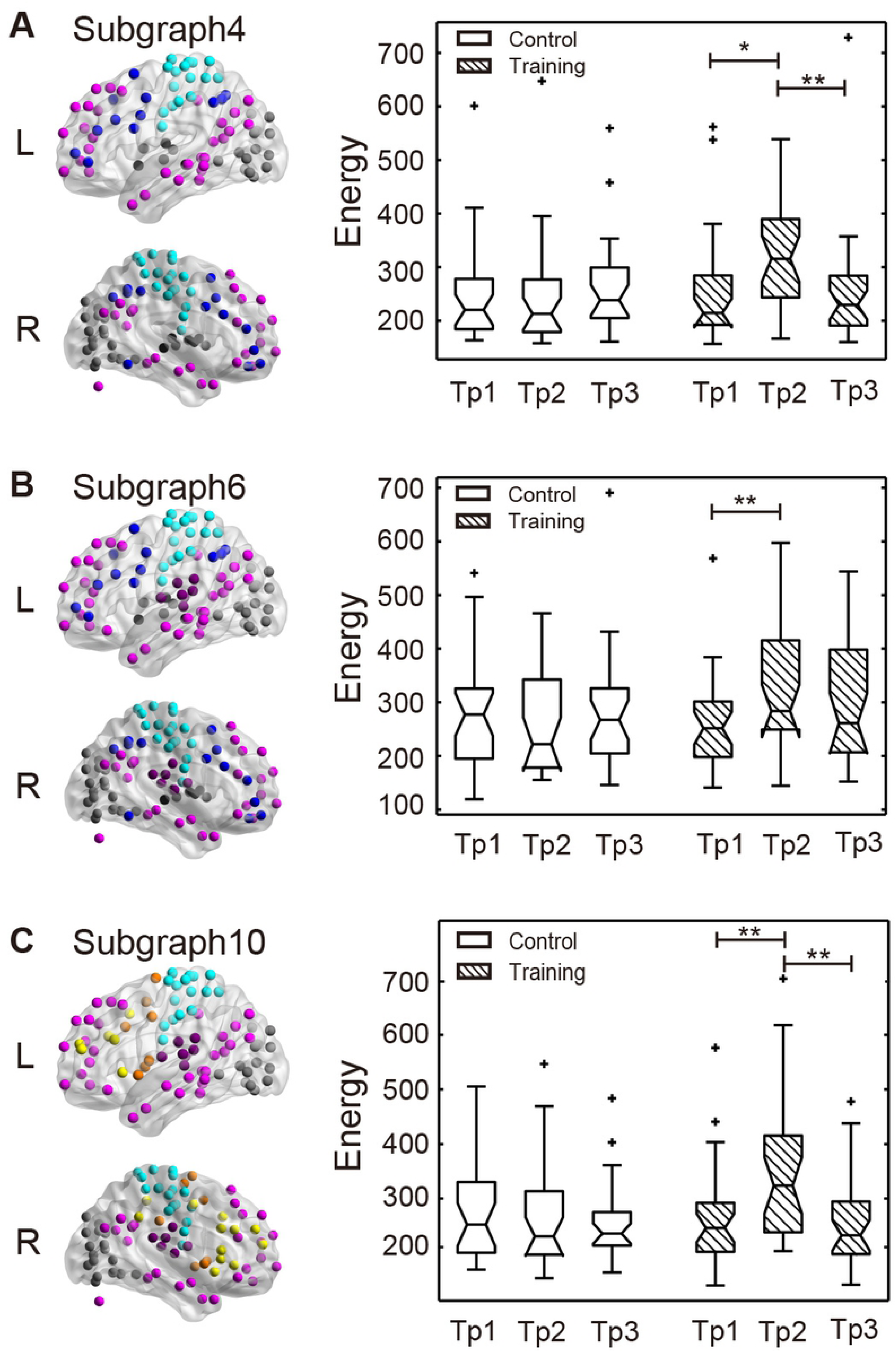
Energy analyses of the time-dependent coefficients. **(A)** Comparison showed increased energy for subgraph 4, including sensory, default mode, visual, and fronto-parietal task control systems significantly expressed, when comparing Tp2 with Tp1 (𝑝 = 0.035), and decreased energy when comparing Tp3 with Tp2 (𝑝 = 0.020) in the training group. **(B)** Comparison showed increased energy for subgraph 6, including sensory, default mode, visual, auditory, and fronto-parietal task control systems significantly expressed, when comparing Tp2 with Tp1 (𝑝 = 0.005) in the training group. **(C)** Comparison showed increased energy for subgraph 10, including sensory, default mode, visual, auditory, cingulo-opercular task control, and salience systems significantly expressed, when comparing Tp2 with Tp1 (𝑝 = 0.006), and decreased energy when comparing Tp3 with Tp2 (𝑝 = 0.003) in the training group. Notes: Tp1, time point 1; Tp 2, time point 2; Tp 3, time point 3.

**Fig 6.**
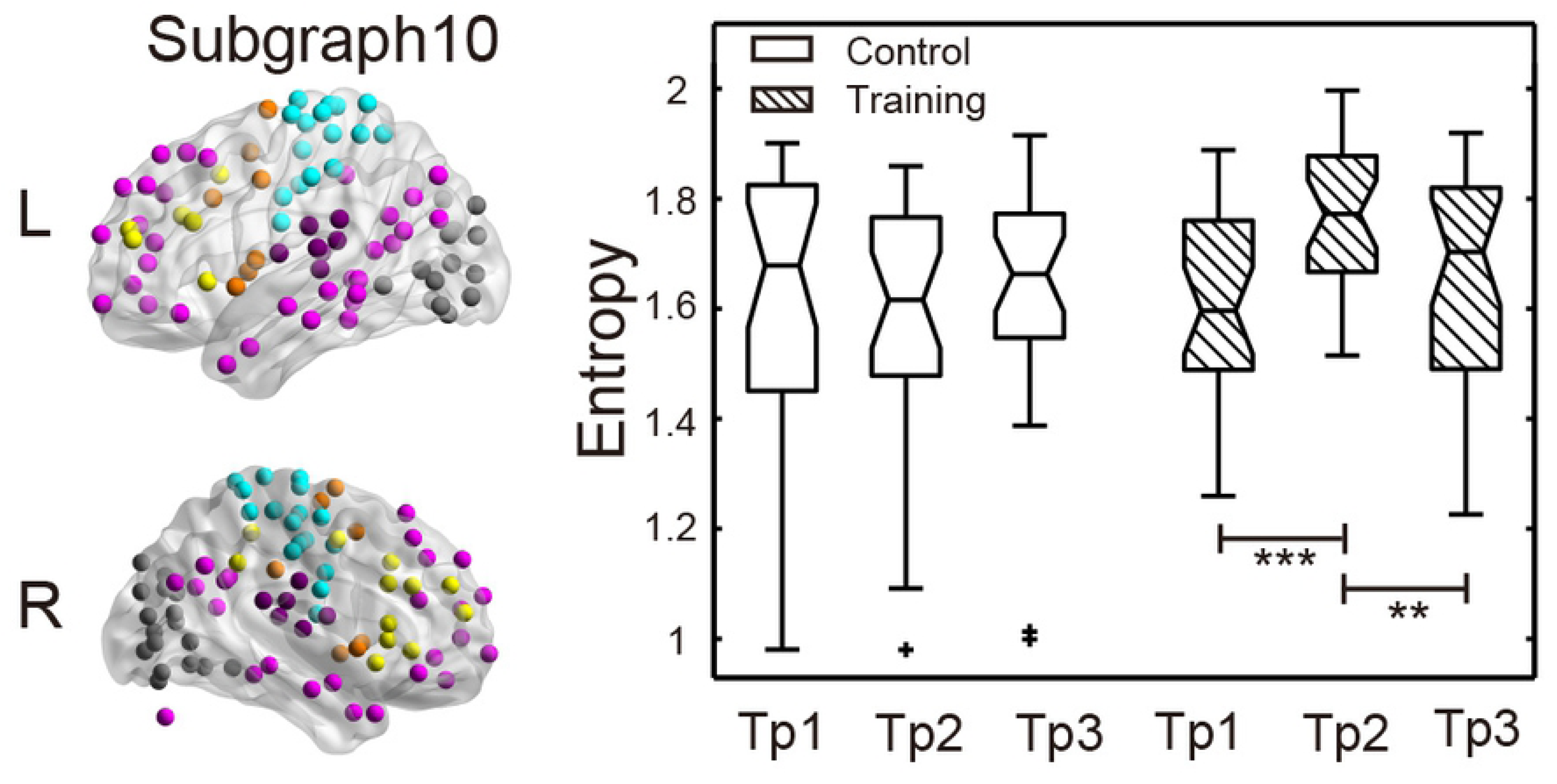
Entropy analyses of the time-dependent coefficients. Subgraph 10 with sensory, default mode, visual, auditory, cingulo-opercular task control, and salience system significantly expressed, showed increased entropy when comparing Tp2 with Tp1 (𝑝 < 0.001), and decreased entropy when comparing Tp3 with Tp2 (𝑝 = 0.001) in the training group. Notes: Tp1, time point 1; Tp 2, time point 2; Tp 3, time point 3.

## Discussion

A large and increasing number of analyses of neuroimaging data have addressed connectivity of functional network in musical training model [23–25]. However, these studies have focused on the static functional connectivity and did not specifically target the nonstationary nature of the functional connectivity that contains a wealth of information. Specifically, our results showed profound and noticeably dynamic interaction of brain functional systems. We also found significant alterations of musical training induced dynamic interaction of brain functional systems.

Nonnegative matrix factorization has been proposed to identify a set of subgraphs of a functional brain network that dynamically vary across subjects and across time, so that the connectivity for a subject in each time window is a nonnegative combination of basis subgraphs [13, 26]. According to its underlying assumptions, functional systems belonging to the same subgraph have a same coactivation pattern and allow interacting with each other. Indeed, multiple studies have shown high correspondence of coactivation pattern and dynamic integration between functional systems from overall functional connectivity analyses [27–29]. Applied to sliding window that offers detecting dynamic functional connectivity, nonnegative matrix factorization effectively probes dynamic interactions of functional systems. Previous analyses in musical training model evaluated static functional connectivity within or between functional systems [23–25]. The current work, on the other hand, targeted the dynamic interaction of functional systems within the assembly of all subjects in a concatenated functional connectivity network. Notably, while our dynamic functional connectivity operated on an unsupervised matrix decomposition method, the obtained nonnegative combination of basis subgraphs and sparse time-dependent expression coefficients eases the neurophysiological interpretability of the relative expression of different subgraphs over time. Our findings, thus, likely reflect dynamic interaction of functional systems in musical training model.

In our study, analyses of association between functional system interaction and musical training induced plasticity showed increased interaction of functional systems in the training group after a period of musical training, particularly with regards to the primary functional systems. Similar results were found in our previous study of static functional connectivity analyses [25], as well as in cross-modal integration studies [30, 31]. Interaction of functional systems increases likely drive networks toward a more collaborative pattern in complex task. A plausible explanation for these changes may be enhanced functional integration after a period of training. The brain works in the form of brain network organization, global integration of local (functionally specialized) interaction, which global integration via long-range weak connections facilitate diverse cognitive function mediated by short-range dense connections [1, 32]. The long-range weak connections are relatively flexible and more likely to be modulated for various functional demands, especially complex tasks involving multiple functional specialized systems. On the other hand, high proportions of cycles and minimized wiring length in structural connection patterns may underlie the functional integration of local interaction, giving rise to the spatially and temporally highly organized functional brain network [32]. These architectures facilitate a high level of information integration [33] and the formation of an integrated ‘dynamic core’ which is the potential neurological system associated with higher cognition and consciousness [34, 35].

As for the primary functional systems, we observed more enhanced interaction between them, however, less for higher-order cognitive-related systems. Considering that musical performance involving interaction of several modalities and higher-order cognitive functions, it is likely that the musical training induced plasticity in young adults may follow a bottom-top mechanism. A large amount of studies has consistently observed plastic effects of musical training on musical related regions [9, 10, 12, 36]. Plastic changes in multisensory integration and strengthening connections between brain regions may also have an effect beyond the most relevant musical related regions, such as the mirror neuron system [37], reward system [38], and subcortical hippocampus [8]. In the light of previous analyses across multiple systems, one may postulate that primary systems are more associated with musical training induced plasticity.

Results from entropy analyses of time-dependent coefficients suggest that interaction between functional systems undergoes significant changes in their fine-scale dynamic after a period of musical training. We observed differences in the time-dependent variability of the subgraph 10, consisting of sensory, visual, auditory, default mode, cingulo-opercular task control and salience systems, after training in the training group when compared with the control group (Fig 6). The changed entropy of time-dependent coefficients in the training group after musical training parallels our previous findings in the dynamic modularity analyses [5], yet with markedly higher effects. Greater switching behaviour of the subgraph illustrates increasing flexibility of the interaction between functional systems after musical training. Noteworthy, the flexible interaction has been shown between functional systems during executive cognition [4], reinforcement learning [39], and musical training [25]. In the current study, flexible interaction was found between the primary functional systems, as well as the higher-order cognitive related systems—cingulo-opercular task control and salience systems.

Piano playing involves sight-read notations and simultaneously producing the appropriate motor action, and then auditory feedback to be checked to identify whether the played notes are well tuned. During the procedure, the primary functional systems, including visual, sensory and auditory are the most relevant and important aspects of music perception [40–42]. Besides, higher-order cognitive related systems for control, adjustment and reward are also critical during this procedure [43, 44]. Findings in the current study suggest the cingulo-opercular task control system and salience system may also play a vital role in piano playing. The cingulo-opercular task control system, a positive-task network, was considered to maintain task goals, sustain adjustments for feedback control, and monitor errors [45, 46]. The salience system is implicated in the detection and integration of emotional and sensory stimuli, as well as in modulating the switch between the internally directed cognition of the default mode regions and externally directed cognition of the central executive network [47]. We can speculate that the cingulo-opercular task control system may work as a monitor and mediator to modulate the information from primary systems and other functional systems during piano performance. Moreover, information of emotional and sensory stimuli was detected and integrated by the salience system, then may transform to default mode regions or higher-order cognitive related systems, so that an accurate and expressed performance could be presented.

It can be seen that the differences between the training group and the control group essentially concern the Tp2 of the measurements, and that at Tp3 the modifications decrease for the training group and become again close to the control group. Alterations were not sustained, which suggested that once the training process was over, the functional networks reversed to their original states. On one hand, unlike the learning induced morphological changes, which may involve slow-evolving mechanisms such as neuronal or glial cell genesis [48], we suggest that the dynamic functional changes resemble fast-adjusting manner. On the other hand, the overall decrease in the changed time-dependent coefficients at the 12 weeks follow-up assessment showed no significant correlation with the practice time. This could be due to the decrease of piano performance at the 12 weeks follow-up. We may speculate the relationship between the decrease of the qualified time-dependent coefficients and the piano performance if it scored.

There are still some limitations in the current study. First, control group in experimental design was only one passive control group without any intervention compared with the training group. It could be better to add an active control group for explanation of observed results [49]. Second, plastic changes of dynamic interaction between functional systems were found by using nonnegative matrix factorization, obtaining a set of subgraphs and their corresponding time-dependent coefficients. The dynamic behaviour was quantized by overall energy and entropy, while we cannot convey how these dynamic interacted functional systems switch their behaviour, and how the switch behaviour further changed by the musical training. Other methods [50] could be used for detecting different aspects of the dynamic switch pattern. Third, each subgraph obtained from nonnegative matrix factorization included more than one functional system, and the corresponding coefficients indicated overall co-expression pattern of functional systems. How these functional systems dynamically changed in the musical training model could be investigated by using sliding window combined with independent component analysis [51]. Future studies could shed more light on these issues.

## Methods

### Longitudinal experiment design

Sixty young adults, all native Chinese speakers, were recruited from local university. All participants meet the criteria: (1) no history of neurologic or psychiatric diseases or health problems affecting dexterity; (2) without any experience of musical training; (3) no depression based on the Beck Depression Inventory (BDI) [52]; (4) right-handers defined by a handedness questionnaire (a modified version of the Edinburgh Handedness Inventory) [53]; (5) the Advanced Measures of Music Audiation (AMMA) score between 20 and 80; (6) IQ score no higher than 140 obtained by the Wechsler Adult Intelligence Scale-Revised Chinese revised version (WAIS-RC) [55].

Participants were randomly enrolled in the training group and control group. Four participants dropped out due to personal reasons during the program, which caused 29 participants (13 males) in the training group and 27 participants in the control group (13 males). Participants in the training group received 24 weeks piano training while the control group without any intervention. The piano training program included professional instructions, practice with the instructions and a final piano performance. Professional musicians provided instructions about music theory, progressive difficulty in piano performance and technical finger motor exercise once a week in the form of one-hour one-to-two teaching sessions. Instructions of music theory and progressive difficulty in piano performance referred to the *Bastien Piano for Adults-Book 1* [56] and technical finger motor exercises referred to the *Hanon Piano Fingering Practice*. A minimum five 30-minute practice sessions (i.e. five days, each day at least 30-minute practice) and maximum seven 60-minute practice sessions (i.e. seven days, each day at most one-hour practice) per week in the assigned room were required after the weekly piano course, and the practice time for each participant was logged. At the end of the training program, participants performed selected pieces from *Bastien Piano for Adults-Book 1* assessed by professional musicians, and those who were able to individually and skilfully complete the selected pieces achieved the equivalent of the Central Conservatory of Music piano level 4.

All participants both in the training group and the control group received behavioural tests and scanning sessions at three time points: at the beginning (Tp1) and the end (Tp2) of 24 weeks training and at 12 weeks after training (Tp3). The repeated behavioural tests at all three time points included three subtests (block design, digit symbol, and digit span) of the Wechsler Adult Intelligence Scale-Revised Chinese revised version and trail making tests (parts A and B). Two sample t-test implemented in SPSS (SPSS version 22) was used to test the demographic data except for the gender (Chi-squared test), musical aptitude test and behavioural tests at baseline. A mixed ANOVA with between-subject factor group (training group and control group) and within-subject factor time (Tp1, Tp2 and Tp3) including age, gender, and education as covariates of no interest was employed to test the interaction effects of group and time on the assessments of the repeated behavioural tests, following a post hoc pairwise t-test between the factor of time.

### Image acquisition

All of the MRI data were obtained using a SIEMENS Trio Tim 3.0T scanner with a 12-channel phased array head coil in the Imaging Centre for Brain Research, Beijing Normal University. The 3D high-resolution brain structural images were acquired using T1-weighted, sagittal 3D magnetization prepared rapid gradient echo (MPRAGE) sequences. The sequence parameters had a repetition time (TR)=2530 ms, echo time (TE)=3.39 ms, inversion time (TI)=1100 ms, flip angle=7°, FOV=256 mm×256 mm, in-plane resolution=256×256, slice thickness=1.33 mm, and 144 sagittal slices covering the whole brain. During the resting state session, the participants were instructed to hold still, stay relaxed and keep their eyes closed but not fall asleep. The functional MRI data were obtained using an echo-planar imaging (EPI) sequence with the following parameters: 33 axial slices, thickness/gap=3.5/0.7 mm, in-plane resolution=64×64, repeat time (TR)=2000 ms, echo time (TE)=30 ms, flip angle =90°, and a field of view (FOV)=200 mm×200 mm. None of the participants fell asleep according to a simple questionnaire after the scan.

### Data preprocessing

Data preprocessing was conducted using the Data Processing Assistant for the Resting-State Toolbox (DPARSF, http://rfmri.org/DPARSF; [57]. The first 10 volumes were discarded allowing for signal equilibrium and adaption for the participants to the circumstances. The remaining data were slice-timing corrected for interleaved and realigned to the first image in the series for head motion correction. Notably, no data were excluded when the head motion excluding criteria were displacement >2.5mm and rotation >2.5°. Then, the resulting data were normalized to the Montreal Neurological Institute (MNI) space and spatial smoothed with an 8-mm Gaussian Kernel. Spurious variances were removed through linear regression, including 24 parameters from head motion correction, the global mean signal, the white matter signal, the cerebrospinal fluid signal, and head motion scrubbing (each “bad” volumes with frame-wise displacement > 0.2 mm and their 1 forward and 2 back neighbours as a regressor) [58]. Finally, the preprocessed data were performed band-pass temporal filtering (0.01-0.1 Hz) to reduce the effect of low frequency drift and high frequency noise [59, 60].

### Dynamic functional network construction

The brain was functionally partitioned into 𝑁 = 264 regions, and for each region, the time series was obtained by averaging the time series of all voxels extracted from denoting 5-mm radius spheres centered on previously reported coordinates [61]. By using sliding time windows, each time series was divided into 𝑇 = 141 time windows with window length 𝐿 = 50 TRs and step of one TR [51, 62], as shown in Fig 1. In each time window, time series was transformed into time-frequency space by using Morlet wavelet transform [63]. The wavelet coherence was used to calculate the functional connectivity between any pair of regions. As shown in Fig 1C, for each subject at each time point (i.e. Tp1, Tp2 and Tp3), we obtained 141 264 × 264 symmetric functional connectivity network in which each element ^𝐴^𝑖𝑗𝑙 represents the wavelet coherence between regions 𝑖 and 𝑗 in time window 𝑙. We extracted the upper triangle of each ^𝐴^𝑖𝑗𝑙 for all subjects at three time points and unfolded each upper triangle matrix in to a vector. Then, all these vectors were concatenated into a matrix 𝑉 with size 𝑁(𝑁 − 1)/2 × 𝑇𝑆, where 𝑆 = 168 is the number of sessions of all subjects at three time points.

### Nonnegative matrix factorization

The sparse nonnegative matrix factorization (NMF) algorithm was employed to decompose the concatenated dynamic functional connectivity matrix to a set of subgraphs and their corresponding time-dependent expression coefficients:

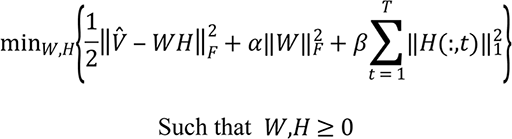

where 𝑉 is the concatenated matrix across time windows, subjects and time points, 𝑊 is a matrix of subgraph connectivity with size 𝑁(𝑁 − 1)/2 × 𝑘, and 𝐻 is a matrix of time-dependent expression coefficients for each subgraph and subject with size 𝑘 × 𝑇𝑆, as shown in Fig 1D. The parameter 𝑘 is the number of subgraph, 𝛽 is the penalty parameter which enforces the sparsity on the expression coefficients matrix and 𝛼 is a parameter which gives an upper on the connection strengths within the functional subgraphs. By performing a grid-search procedure over a range of parameters 𝑘 and 𝛽, the residual sum of squares, defined as 𝑅𝑆𝑆 = ‖𝑉 − 𝑊𝐻‖_𝐹_, was computed to optimize the values for hyperparameters 𝑘 and 𝛽.

The 264 nodes were mapped to 13 well-known functional systems, including sensory, cingulo-opercular task control, auditory, default mode, memory retrieval, visual, fronto-parietal task control, salience, subcortical, ventral attention, dorsal attention, cerebellar and uncertain systems [61]. For each obtained subgraph, 1000 permutation tests were performed to determine the most highly expressed functional systems. Functional system labels were randomly permuted 1000 times and the mean connections in each system was computed to construct the null distribution. Significantly expressed functional system was defined if its connection strength was above the 95% confidence interval threshold.

To quantify the subgraph’s dynamic changes in expression, the signal energy of the time-dependent coefficients for each subgraph and subject was computed, which was defined as 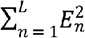, where 𝐸_𝑛_ is time-dependent coefficients for each subgraph and subject in each time window and 𝐿 is the length of the signal for a subject. To quantify the dynamic switching behaviour of subgraph expression, the signal entropy was computed using a histogram-based entropy estimator method that computed the entropy 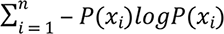, where 𝑃(𝑥) is a probability mass function of the time-dependent coefficients computed using the histogram [64].

### Statistical analysis

A two-sample *t* test for demographic data and a Chi-squared test for gender were performed to test whether there were any significant differences between the two groups at baseline. A mixed ANOVA with a between-subject factor group (training group and control group) and within-subject factor time (Tp1, Tp2 and Tp3) was used to test the interaction effect of the group and time on the repeated measured behavioural tests with age, gender and education as covariates of no interest. Significant interactions were followed by a Bonferroni post hoc pairwise *t*-test between the factor of time to determine which of the time points differ from each other.

For each subgraph, the mixed ANOVA with a between-subject factor group (training group and control group) and within-subject factor time (Tp1, Tp2, and Tp3) was employed to test the interaction effect of the group and time on the dynamic signal energy and entropy with age, gender and education as covariates of no interest. Significant interactions were followed by a Bonferroni post hoc pairwise *t*-test between the factor of time. In addition, correlation analysis was performed between the subgraph’s dynamic changes (only those for which the post hoc pairwise t test was significant) and practice time in the training group.

### Ethics Statement

The Bioethics Committee of Beihang University approved the study. The written informed consent was obtained from each participant with a complete description of the study.

## Acknowledgements

We gratefully acknowledge the assistance of musicians Xia Zhao and Jie Deng for their professional instructions on the musical training programme and all volunteers for their continuing participations in this longitudinal study.

